# Metabolic benefits of dietary medium-chain fatty acids bypass hepatic ketogenesis

**DOI:** 10.1101/2025.10.08.681190

**Authors:** Ye Cao, Kyle Feola, Wenke Jonas, Pascal Gottmann, Stephanie K. Holm, Ricardo Monroy, Josephine M. Kanta, Kasper Suhr Jørgensen, Christopher A. Bishop, Christoffer Clemmensen, Annette Schürmann, Andreas M. Fritzen, Sarah C. Huen, Maximilian Kleinert

## Abstract

Saturated medium-chain fatty acids (MCFAs) are ketogenic nutrients, and their dietary intake has been linked to health benefits such as improved energy balance and increased insulin sensitivity. Here, we tested the hypothesis that MCFA-mediated hyperketonemia is required for these benefits. Using tissue-specific 3-hydroxy-3-methylglutaryl-CoA synthase 2 (*Hmgcs2*) knockout mouse models, we show that the hyperketonemia induced by oral administration of triacylglycerols with the MCFA octanoic acid (C8:0-MCT) is mediated by hepatic, but not intestinal, ketogenesis. Remarkably, acute reductions in food intake and body weight induced by C8:0-MCT in obese mice were independent of hepatic ketogenesis; likewise, chronic protection against weight gain and marked improvements in insulin sensitivity elicited by MCFA-rich diets did not require hepatic ketogenesis. These data demonstrate that dietary MCFAs are potent bioactive lipids capable of inducing metabolic benefits independently of their ketogenic properties.

## Introduction

Dietary saturated medium-chain fatty acids (MCFAs) have emerged as promising nutritional interventions for metabolic disorders. They differ from long-chain fatty acids (LCFAs) in absorption, transportation, and metabolic handling, and have been linked to improved glycemic control, increased postprandial energy expenditure, and body weight regulation in mice and humans^1^. Despite longstanding research interest and expanding clinical applications, the mechanisms by which MCFAs improve metabolic health remain elusive. One distinctive feature of MCFA metabolism is the rapid induction of nutritional hyperketonemia, which underlies the wide use of MCFAs as components of ketogenic diets^2–5^. Beyond serving as alternative fuels during carbohydrate deficiency, ketone bodies have emerged as signaling molecules that regulate multiple aspects of systemic metabolism, including glucose homeostasis, substrate utilization, hormone secretion, and inter-organ communication^6,7^. Consequently, the potent ketogenic effect of MCFAs has led to the widespread assumption that many of their metabolic benefits are substantially mediated by ketone body production^8,9^. However, whether ketogenesis is causally required for the metabolic actions of dietary MCFAs has not been directly tested.

The tissue source of nutritional hyperketonemia elicited by ingestion of MCFAs also remains unresolved. Although the liver is considered the major ketogenic organ and is found to be solely responsible for fasting-induced ketogenesis^10,11^, the rate-limiting enzyme for ketogenesis, mitochondrial 3-hydroxy-3-methylglutaryl-CoA synthase 2 (HMGCS2), is also expressed in the intestine^12^, where ingested nutrients first pass through. In the small intestine, HMGCS2 has been shown to be required for local ketone body production, with a direct impact on intestinal stem cell homeostasis^12^, raising the possibility that both the liver and the intestine contribute to nutritional ketogenesis following MCFA ingestion. Therefore, defining the tissue source of MCFA-mediated nutritional hyperketonemia is a necessary prerequisite for addressing the role of ketogenesis in MCFA actions.

Here, using constitutive liver- and intestine-specific *Hmgcs2* knockout mice, we identified the liver as the primary source of MCFA-induced ketogenesis following oral administration of the highly ketogenic medium-chain triacylglycerol (MCT) C8:0-MCT^13–15^. Next, we tested whether hepatic ketogenesis mediates the acute and chronic metabolic responses to dietary MCFAs, including reduced food intake and improved insulin sensitivity, and found that these effects occur independently of hepatic ketogenesis.

## Result

### C8:0-MCT acutely elevates ketone body levels in portal vein blood

Given that *Hmgcs2* is expressed in the intestine (Fig. 1a), we first explored whether the intestine contributes to MCFA-mediated hyperketonemia by assessing β-hydroxybutyrate (β-OHB) levels in blood from the portal vein, which transports blood from the intestine to the liver, and compared them to the systemic blood concentrations following oral administration of C8:0-MCT in lean, chow-fed male mice (Fig. 1b). C8:0-MCT increased circulating β-OHB levels in the systemic blood from 0.60 mmol/L to 2.23 mmol/L (*P* < 0.0001), whereas long-chain triacylglycerol (LCT, olive oil) or water had no effect on circulating β-OHB in the systemic blood at one hour post oral administration (Fig. 1c). Interestingly, with C8:0-MCT, circulating β-OHB concentrations were ~60% higher in portal compared to systemic blood (3.59 mmol/L vs. 2.23 mmol/L, *P* < 0.0001; Fig. 1c), suggesting that the intestine may contribute to the elevated systemic levels of β-OHB.

**Fig. 1.**
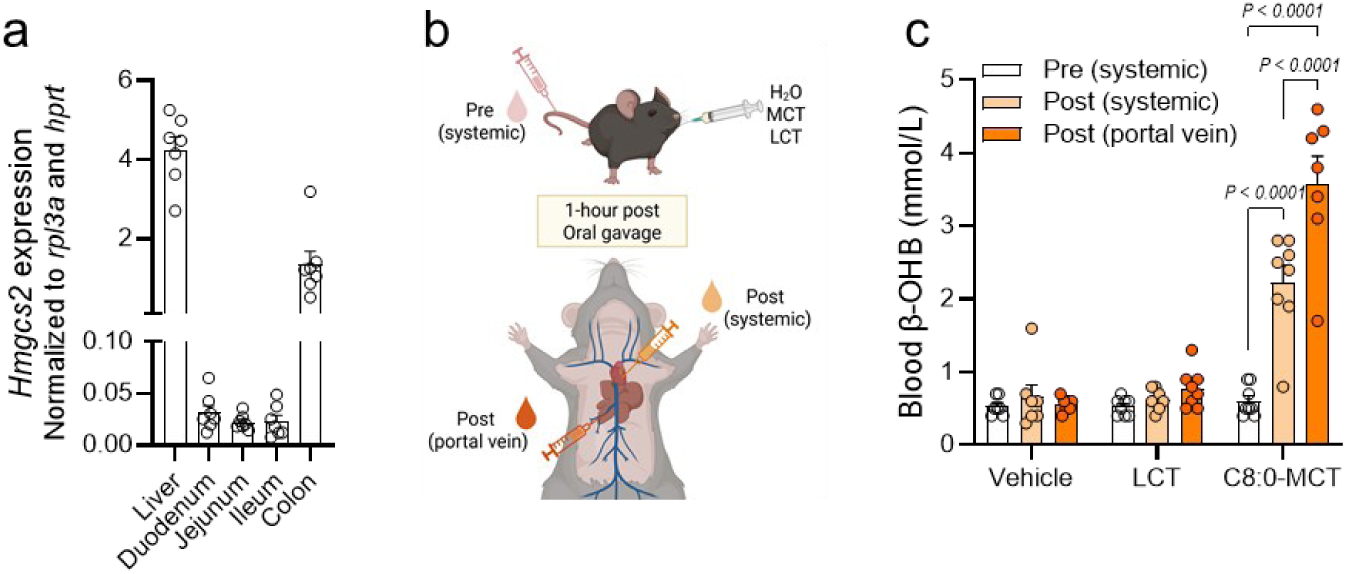
Dietary medium-chain triacylglycerol (MCT) induces potentiated portal vein ketone body concentrations compared with systemic blood. a, mRNA abundance of *3-hydroxy-3-methylglutaryl-CoA synthase 2* (*Hmgcs2*) in the liver, duodenum, jejunum, ileum, and colon from chow-fed male C57BL/6J mice, n = 7. b, schematic overview of the experimental setup in measuring β-hydroxybutyrate (β-OHB) levels in (c) systemic circulation before and at one hour after oral administration, and in portal vein blood after oral administration of water (vehicle), long-chain triacylglycerol (LCT) oil (olive oil) or C8:0-MCT oil at 5 µL/g body weight, n = 7-8. Statistical significance was determined by two-way ANOVA with Tukey’s post hoc test. Data are shown as mean + SEM. Schematic overview created by BioRender.com.

### Hepatic, but not intestinal, ketogenesis mediates C8:0-MCT-induced nutritional hyperketonemia

To directly test the intestinal contribution to MCFA-induced ketogenesis, we generated a mouse model with intestinal *Hmgcs2* deletion: *Villin-Cre*;*Hmgcs2*^fl/fl^ (or intestinal *Hmgcs2* knockout, *Hmgcs2* iKO, Fig. 2a, Fig. S1), and confirmed the deletion by assessing HMGCS2 protein in the intestinal tissues (Fig. S1c-f, i-n). HMGCS2 protein abundance showed a slight (−28%) but statistically significant reduction in the liver (Fig. S1a, g), while it remained unaltered in the kidney of *Hmgcs2* iKO mice (Fig. S1b, h). In lean, chow-fed wild-type mice (*Hmgcs2* WT), systemic β-OHB levels increased after C8:0-MCT gavage, reaching 1.34 mmol/L and 1.57 mmol/L in males and females, respectively, at one hour post-administration, a threefold increase over the baseline, whereas LCT demonstrated no ketogenic effect (Fig. 2b, c). However, in *Hmgcs2* iKO mice, β-OHB levels also increased to the same extent with gavage of C8:0-MCT, indicating that intestinal *Hmgcs2* is dispensable for C8:0-MCT induced hyperketonemia.

**Fig. 2.**
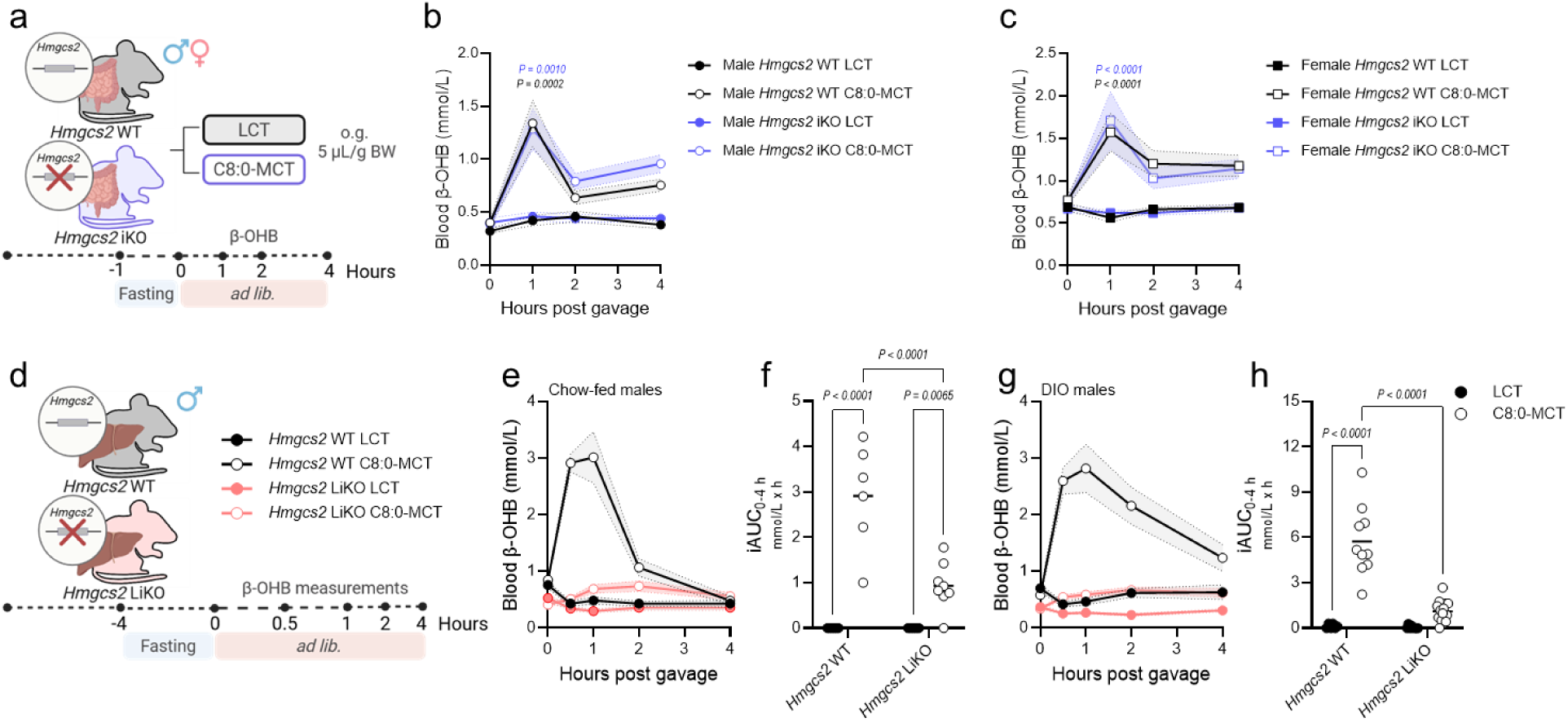
Hepatic, but not intestinal 3-hydroxy-3-methylglutaryl-CoA synthase 2 (HMGCS2) contributes to dietary medium-chain triacylglycerol (MCT)-induced hyperketonemia. a, experimental design for ketone dynamics after oral administration of C8:0-MCT or LCT at 5 µL/g body weight per mouse in *Hmgcs2* WT or *Hmgcs2* iKO mice. b-c, circulating β-hydroxybutyrate (β-OHB) levels before and after oral administration of C8:0-MCT or LCT (olive oil) at indicated time points in male (b) and female (c) *Hmgcs2* WT or *Hmgcs2* iKO mice, n = 5-14 (male), n = 8-11 (female). d, experimental design for ketone dynamics after oral administration of C8:0-MCT or LCT in *Hmgcs2* WT or *Hmgcs2* LiKO mouse. d-h, chow-fed (e, f), or diet-induced obese (DIO, g, h) male C57BL/6J mice were gavaged with C8:0-MCT or LCT (lard) control oil at 3 µL/g (e, f, n = 6-8) or 4 µL/g (g, h, n = 6-16) body weight, followed by blood β-hydroxybutyrate (β-OHB) measurements at 0, 0.5, 1, 2, and 4 h (f, h), with incremental area under the curve (iAUC) (f, h). Diet-induced obesity was induced by feeding young adult mice on a high-fat, high-sucrose diet (HFHSD, S8022-E188, modified from D12492). Statistical significance was determined by two-way ANOVA with Šídák’s post hoc test (b, c) or with Fisher’s LSD (f, h). Data are shown as mean + SEM. Experimental design created by BioRender.com.

Before undergoing β-oxidation, MCFAs must first be activated to their corresponding acyl-CoAs by medium-chain acyl-CoA synthetases (ACSMs) (Fig. S2a). Consistent with the liver being a major site of MCFA oxidation^16^, *Acsm1*, *Acsm3*, and *Acsm5* are highly abundant in this tissue and contribute to MCFA activation^17^. In line with this, we observed detectable mRNA expressions of *Acsm1* and *Acsm5* in the livers of chow-fed mice (Fig. S2b, d). *Acsm3* mRNA was enriched in the kidney (Fig. S2c), another organ capable of oxidizing MCFAs^16^. Isoforms of *Acsm* were at or below the detection limit in intestinal tissues (Fig. S2b-d), and we observed similarly low levels in publicly available transcriptomic datasets from mouse^18^ and human^19^ small intestine (Fig. S2e-g): While *Hmgcs2/HMGCS2* was expressed in small intestinal cells across species and increased with high-fat diet (HFD) feeding in mice or obesity in humans, expression of the different *Acsms/ACSMs* was negligible across intestinal epithelial cell types, even under obese conditions. To directly assess the capacity of intestinal cells to generate ketone bodies from MCFA, we treated human and mouse intestinal cells with C8:0 in vitro. Neither cell type showed evidence of induced ketogenesis, whereas C8:0 increased β-OHB production approximately threefold in murine primary hepatocytes compared with the LCFA control palmitic acid (C16:0) (Fig. S2h). Together, these findings suggest that limited MCFA activation may contribute to the low capacity of the intestine for C8:0-mediated ketogenesis.

Since intestinal *Hmgcs2* is dispensable for C8:0-MCT-induced hyperketonemia, we generated a mouse model with hepatic *Hmgcs2* deletion (*Alb-Cre^+/-^*;*Hmgcs2*^fl/fl^, liver *Hmgcs2* knockout, *Hmgcs2* LiKO) to directly test the role of hepatic *Hmgcs2* in MCFA-induced hyperketonemia (Fig 2d). While oral gavage with C8:0-MCT increased circulating β-OHB in male and female *Hmgcs2* WT mice, hepatic *Hmgcs2* deletion largely abolished C8:0-MCT-induced hyperketonemia in both chow-fed and DIO *Hmgcs2* LiKO mice (Fig. 2e-h, Fig. S3c, d). These data indicate that C8:0-MCT-induced hyperketonemia is chiefly mediated by the liver in both lean and obese mice.

### Hepatic ketogenesis is not required for C8:0-MCT-induced food intake and body weight-lowering effects in diet-induced obese mice

With the liver as the primary source of C8:0-MCT-driven hyperketonemia, we used the *Hmgcs2* LiKO model to determine whether hepatic ketogenesis is required for C8:0-MCT-associated acute metabolic responses (Fig. 3a). In line with our previous findings^20^, C8:0-MCT ingestion reduced food intake by ~28% (diet effect *P* = 0.0159) and promoted body weight loss (MCT −0.46% vs. LCT 0.11%, diet effect *P* = 0.0399) over 24 h relative to LCT in DIO *Hmgcs2* WT mice (Fig 3b-d). Both effects were maintained in *Hmgcs2* LiKO mice, suggesting that hepatic ketogenesis is not required for C8:0-MCT-mediated anorectic and body-weight-regulating responses. We have shown that the C8:0-MCT-associated reduction in food intake is partially mediated by growth differentiation factor 15 (GDF15)^20^. At an hour post administration, C8:0-MCT increased circulating GDF15 levels to 1214.01 pg/mL in *Hmgcs2* WT mice, a threefold increase compared to LCT (418.24 pg/mL) (Fig. 3e). Notably, C8:0-MCT intake induced more pronounced GDF15 secretion in *Hmgcs2* LiKO mice, at 2504.89 pg/mL, which was over fourfold higher than that with LCT (538.74 pg/mL), suggesting that loss of hepatic ketogenesis may enhance selected endocrine adaptations to C8:0-MCT intake, although this did not further decrease food intake. Locomotor activity, energy expenditure, and whole body fatty acid oxidation were unaffected by hepatic *Hmgcs2* deletion or C8:0-MCT administration (Fig. 3f, Fig. S3e, f, i, j), suggesting that the body-weight lowering effects are primarily driven by decreases in energy intake. Respiratory exchange ratio (RER) and whole body carbohydrate oxidation were lower in *Hmgcs2* LiKO mice following administration of either C8:0-MCT or LCT, indicating that hepatic ketogenesis broadly influences postprandial carbohydrate utilization (Fig. S3g, h, k, l). Overall, acute C8:0-MCT ingestion was associated with reduced food intake and body weight, and increased circulating GDF15 levels in male DIO mice, but hepatic ketogenesis is dispensable for these effects.

**Fig. 3.**
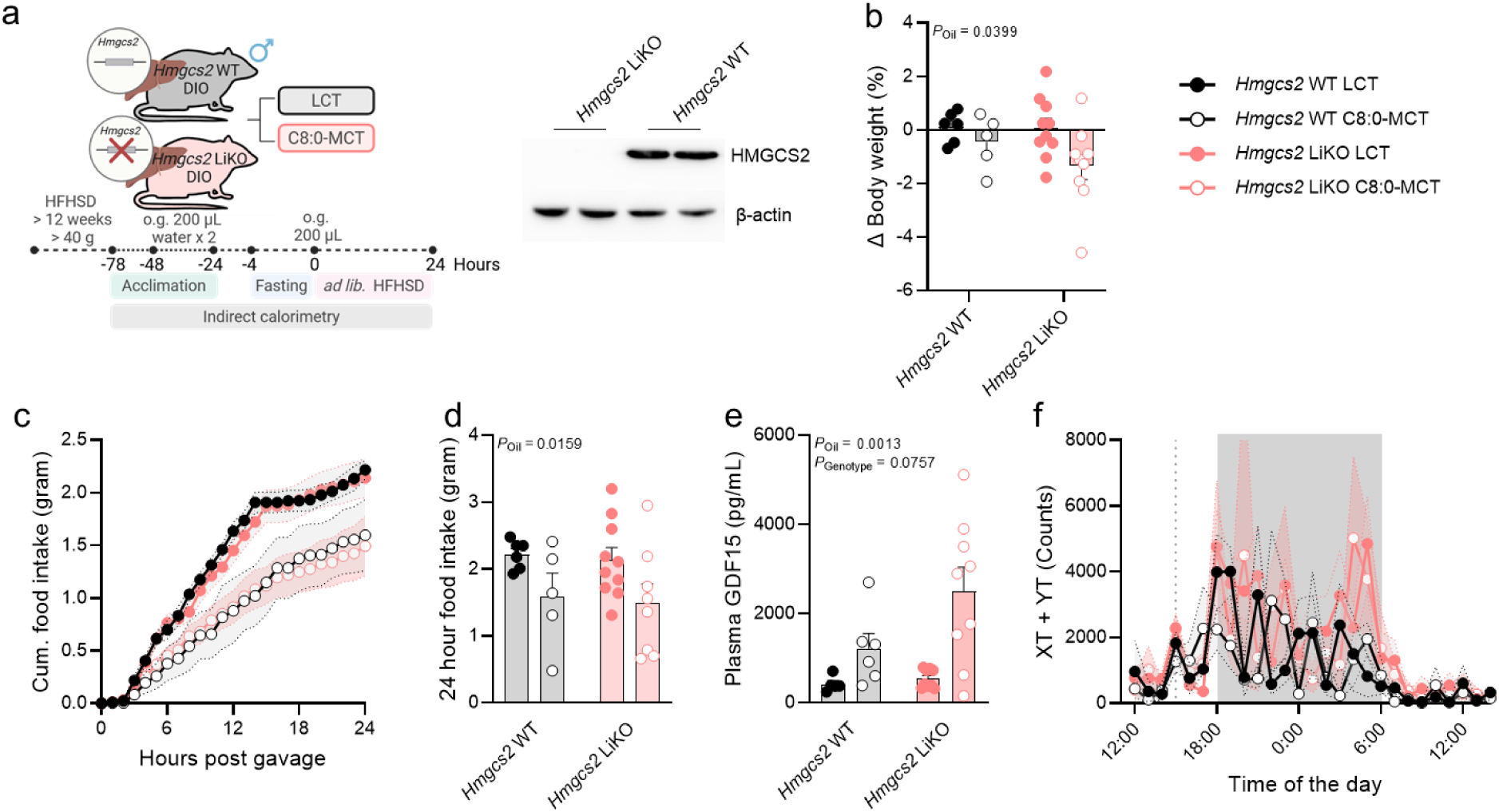
Hepatic 3-hydroxy-3-methylglutaryl-CoA synthase 2 (HMGCS2) is not required for acute MCFA effects on body weight and the regulation of food intake. a, experimental design for metabolic phenotyping in indirect calorimetry cages after oral administration of C8:0-MCT or LCT at 150 µL per mouse in *Hmgcs2* WT or *Hmgcs2* LiKO mice, n = 5-10. Diet-induced obesity was induced by feeding young adult mice on a high-fat, high-sucrose diet (HFHSD, S8022-E188, modified from D12492). b, percent change in body weight at 24 h post gavage. c, cumulative food intake over 24 h. d, 24 h food intake over 24 h. e, plasma GDF15 concentration measured at 1 h after gavage of 4 µL/g body weight of C8:0-MCT or LCT, n = 5-9. f, locomotor activity, shown as combined x + y total movements over 24 h. Statistical significance was determined by two-way ANOVA. Data are shown as mean +/- SEM (c, f) or individual points with mean + SEM (b, d, e). Experimental design created by BioRender.com.

### MCFA-enriched HFD prevents body weight gain and improves insulin sensitivity independent of hepatic ketogenesis

We previously showed that replacing dietary LCFAs with MCFAs C8:0 and C10:0 protects against weight gain and excessive energy intake during HFD feeding^21^. We therefore asked whether hepatic ketogenesis is required for these benefits of MCFAs. *Hmgcs2* WT and *Hmgcs2* LiKO mice were fed a chow, a 45 kcal% LCFA-rich HFD (LCFA HFD), or an isocaloric 45 kcal% C8:0- and C10:0-rich HFD (MCFA HFD) for 6 weeks (Fig. 4a). MCFA HFD-fed mice were protected from weight gain irrespective of hepatic *Hmgcs2* expression, whereas LCFA HFD feeding increased body weight by 15.36% in WT and 20.84% in LiKO mice over 42 days (Fig. 4b, c; Fig. S4a). MCFA HFD also reduced daily energy intake by 37% relative to the LCFA HFD and by 22% relative to chow (Fig. 4d; Fig. S4b). The lower body weight of MCFA HFD-fed mice was largely driven by reduced fat mass (Fig. 4e; Fig. S4d). Lean mass, although increased significantly with chow and LCFA HFD feeding, was broadly comparable among dietary groups (Fig. 4e; Fig. S4c-e). Importantly, these effects were preserved in *Hmgcs2* LiKO mice, demonstrating that hepatic ketogenesis is dispensable for the protection against hyperphagia, weight gain, and adiposity conferred by chronic MCFA intake.

**Fig. 4.**
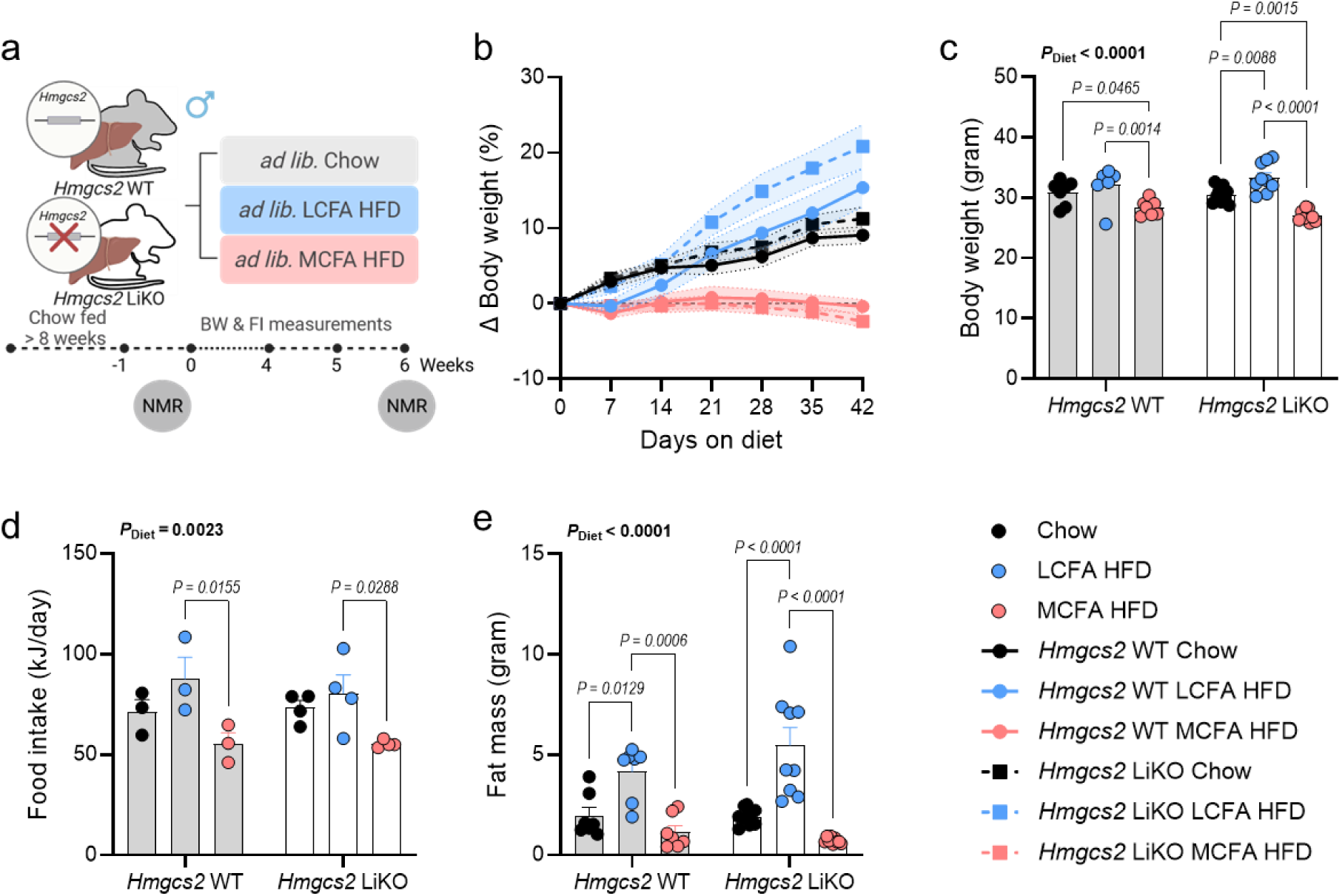
Dietary MCFAs reduce energy intake and body weight independent of hepatic ketogenesis. a, experimental design for chronic metabolic phenotyping following a 6-week dietary intervention of chow, LCFA HFD, or MCFA HFD in *Hmgcs2* WT or *Hmgcs2* LiKO mice, n = 7-9. b-e, weekly body weight change (b), terminal body weight (c), daily energy intake (d), and terminal fat mass (e) determined in mice fed indicated diets over 6 weeks. Statistical significance was determined by two-way ANOVA with Tukey’s post-hoc test. Data are shown as mean +/- SEM (b) or individual points with mean + SEM (c-e). Experimental design created by BioRender.com.

As previously observed^21^, MCFA HFD-fed mice displayed markedly improved glucose tolerance compared with LCFA HFD-fed mice, with glucose excursions comparable to those of chow-fed mice (Fig. 5a-c; diet effect, *P* < 0.0001). Fasting plasma insulin concentrations were more than 70% lower in MCFA HFD-fed mice than in LCFA HFD-fed mice and approximately 50% lower than in chow-fed mice, irrespective of genotype (Fig. 5d). Consistent with this, the surrogate marker of insulin sensitivity in the fasted state, HOMA-IR was strongly affected by diet (*P* < 0.0001) (Fig. 5e). LCFA HFD feeding doubled HOMA-IR relative to chow in WT mice, whereas MCFA HFD feeding reduced HOMA-IR by approximately 80% compared with LCFA HFD and by 60% compared to chow. This improvement was fully retained in *Hmgcs2* LiKO mice. Moreover, glucose-stimulated insulin secretion was substantially lower following MCFA HFD feeding, with the 15-min increment in circulating insulin more than twofold higher after LCFA HFD feeding in both WT and LiKO mice (Fig. 5f, g; Fig. S4f). Thus, dietary MCFAs markedly reduce the insulin requirement to maintain and restore glucose homeostasis, and this improvement in whole-body insulin sensitivity does not require hepatic ketogenesis.

**Fig. 5.**
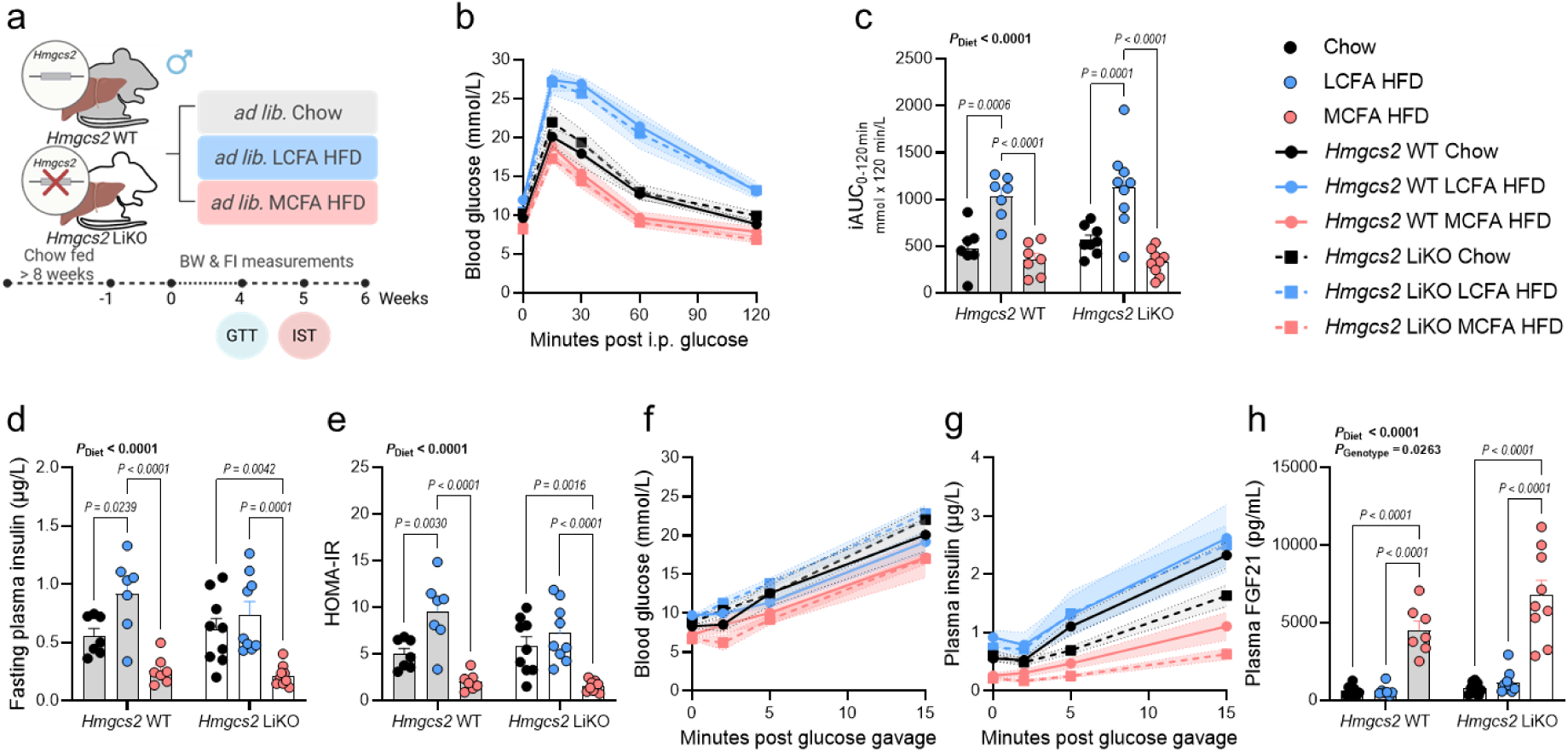
Dietary MCFAs improve glycemia and insulin dynamics independent of hepatic ketogenesis. a, experimental design for chronic metabolic phenotyping following a 6-week dietary intervention of chow, LCFA HFD, or MCFA HFD in *Hmgcs2* WT or *Hmgcs2* LiKO mice, n = 7-9. b, c, glucose excursion curves (b) and the incremental area under these curves (iAUC) (c) determined during an i.p. glucose tolerance test (GTT, 2 g of glucose per kg body weight) conducted 4 weeks after placing mice on the indicated diets. d-g, fasting plasma insulin (d), HOMA-IR (e), glucose excursion curves (f), insulin excursion curves (g, n = 6-9) determined during an oral glucose-stimulated insulin (IST) secretion test (3 g of glucose per kg body weight) conducted 5 weeks after placing mice on the indicated diets. h, plasma FGF21 concentrations at 6 weeks after putting mice on indicated diets. Statistical significance was determined by two-way ANOVA with Tukey’s post-hoc test. Data are shown as mean +/- SEM (b, f, g) or individual points with mean + SEM (c-e, h). Experimental design created by BioRender.com.

Finally, we examined circulating fibroblast growth factor 21 (FGF21), a hepatokine with established effects on systemic metabolism^22^, which we have previously shown to be induced by chronic MCFA intake^21^. After 6 weeks, circulating FGF21 concentrations were approximately sevenfold higher in MCFA HFD-fed mice than in chow- or LCFA HFD-fed mice (Fig. 5h; diet effect, *P* < 0.0001). Similar responses were observed in *Hmgcs2* LiKO mice, which displayed overall higher circulating FGF21 levels across all dietary conditions. Collectively, the protection against hyperphagia, weight gain, and adiposity; marked improvements in glucose homeostasis and insulin sensitivity; and robust induction of FGF21 persisted in the absence of hepatic ketogenesis, demonstrating that hepatic ketogenesis is dispensable for the chronic metabolic benefits of dietary MCFAs.

## Discussion

MCFAs induce rapid ketogenesis, which has been proposed to be a key mediator of their metabolic benefits. Here, we identify the liver as the primary source of C8:0-MCT-induced nutritional hyperketonemia and demonstrate that hepatic ketogenesis is dispensable for many of the acute and chronic metabolic benefits of intake of MCFAs, including regulation of energy intake, body weight, adiposity, endocrine responses, glucose homeostasis, and insulin sensitivity.

We first clarified the tissue source of MCFA-induced hyperketonemia. C8:0-MCT administration resulted in a greater increase in ketone body levels in portal vein blood than in the systemic circulation, initially suggesting intestinal contribution to nutritional hyperketonemia, in line with prior work proposing intestinal contribution to circulating ketone bodies on an HFD^23^. However, intestinal-specific *Hmgcs2* deletion did not block C8:0-MCT-induced hyperketonemia, whereas hepatic deletion abolished it, indicating that the liver, not the intestine, is the central organ for nutritional hyperketonemia. These findings reaffirm previous observations that C8:0-MCT is highly ketogenic in humans and mice^13–15,20^, and extend existing knowledge that hepatic metabolism is the indispensable driver of C8:0-MCT-mediated hyperketonemia.

Notably, these findings do not exclude intestinal ketogenesis from other substrates, as acyl-CoA synthetases for other fatty acids are highly expressed in intestinal regions^24^. The short-chain fatty acid butyrate (C4:0) has been shown to induce ketogenesis in cultured human colonic cancer epithelial cells and jejunal enteroids, partially dependent on HMGCS2^25^. In mice, ketogenic diet feeding has been shown to elevate ketone body levels by approximately sevenfold in small intestinal crypts, accompanied by increased HMGCS2 expression^12^. In contrast, intestinal cells failed to generate ketone bodies from C8:0 in vitro, coinciding with low expression of several medium-chain acyl-CoA synthetases to activate MCFAs. These findings suggest that intestinal ketogenesis is likely substrate-dependent, relying on enzymes to activate substrates, and highlight that examining HMGCS2 expression alone is insufficient to determine whether a given cell type can produce ketone bodies from a specific energy source.

Acute and chronic metabolic effects of MCFAs were largely preserved in mice lacking hepatic *Hmgcs2*, indicating that alternative pathways, other than ketogenesis, mediate these actions. We have previously identified substantial liver transcriptomic remodeling following acute and chronic MCFA intake, with GDF15 and FGF21 emerging as prominent MCFA-regulated endocrine signals^20,21^. GDF15 provides a direct mechanism for hypophagia, as it is required for the full anorectic effect of C8:0-MCT^20^. The mechanisms underlying favorable glucose and insulin homeostasis are less well known. MCFAs protect against HFD-induced insulin resistance in healthy individuals^26^, and prevent HFD-induced hyperinsulinemia and impaired glycemia in mice^21^. Although MCFAs activate hepatic cyclic adenosine monophosphate (cAMP)-responsive element-binding protein H (CREBH)-FGF21 signaling, they improve glucose homeostasis and lower insulin demand independent of *Fgf21*^21^, and, here, of hepatic *Hmgcs2*, indicating these effects may involve other MCFA-responsive signatures^15,20,21^. For instance, this could be attributed to the rapid activation of nutrient-sensing pathways, such as the peroxisome proliferator-activated receptor (PPAR) signaling, following the intake of natural ligands MCFAs^27–29^, as the insulin-sensitizing effect of PPAR activation does not require FGF21^30^. However, the contribution of PPAR signaling or other MCFA regulatory pathways remains to be examined directly.

More broadly, our study resolves a paradox in MCFA metabolism and adds to a growing body of work challenging the assumptions that ketogenesis is required for several adaptations historically linked to an increase in circulating ketone bodies, including starvation^11^, caloric restriction^31^, exercise training-mitigated liver steatosis^32^, and ketogenic diet-mediated intestinal stem cell activity and intestinal tumorigenesis^33^. Supporting this notion, we have recently shown that closely related MCFAs with similar ketogenic capacity, C6:0 and C8:0, engage markedly distinct physiological and molecular programs, including energy intake, endocrine response, fuel selection, hepatic transcription, and circulating metabolome^15^. Together, these findings argue against using hyperketonemia as a surrogate for the metabolic actions of MCFAs, and mechanisms beyond ketogenesis need to be considered, especially since it was recently shown that MCFA intake regulates hundreds of circulating metabolites^15^.

Hepatic *Hmgcs2* deletion nevertheless induced physiological adaptations independent of MCFA-specific responses. For the first time, we show that hepatic ketogenesis contributes to acute systemic substrate utilization in response to lipid exposure, as loss of hepatic ketogenesis decreased postprandial RER and carbohydrate oxidation following both C8:0-MCT and LCT administration. Lowered RER has also been observed in female global *Hmgcs2* KO mice during caloric restriction, in which post-feeding increased RER was blunted^31^, supporting a broader role for ketogenesis in substrate selection. Ketogenesis generally serves as an important metabolic outlet for MCFA-CoA-derived acetyl-CoA, contributing to mitochondrial homeostasis^6^. When this route is restricted, acetyl-CoA must be redirected to alternative fates. Impaired ketogenesis has been linked to overloaded tricarboxylic acid (TCA) cycle flux and substantially altered glycogen, glucose, and lipid metabolism^34,35^, which may explain the alterations in fuel partitioning observed here and warrant future studies.

Our study has several limitations. Hepatic *Hmgcs2* deletion eliminates peripheral C8:0-MCT-induced hyperketonemia, but local ketogenesis in other tissues, such as the brain, heart, and kidney^11,36,37^, should not be overlooked. We addressed only a subset of metabolic effects following selective MCFA intake, and ketone bodies may still contribute to responses in other tissues, including improved brain function^8,38–41^, where they can serve as oxidative substrates and signaling metabolites^7^. We identified hepatic ketogenesis as dispensable for the observed MCFA effects, but the mechanisms linking loss of ketogenesis to enhanced endocrine signaling and altered fuel selection remain inferential, and the downstream ketone-independent mechanisms are unresolved. In addition, acute and chronic metabolic phenotyping were primarily conducted in young adult male mice, and we cannot rule out that hepatic ketogenesis does not contribute to MCFA-mediated metabolic effects in females, or across the lifespan.

In summary, hepatic ketogenesis is required for C8:0-MCT-induced nutritional hyperketonemia but is dispensable for the major acute and chronic metabolic benefits of dietary MCFAs. Thus, the assumption that the mechanisms underlying ketogenesis largely contribute to MCFA function may be inappropriate, and these lipids are bioactive in a much broader sense than their potent ketogenic properties.

## Method

### Animals

Male C57BL/6J mice (Charles River, Germany, or Janvier, France), female and male (*Hmgcs2*^fl/fl^) wild-type (*Hmgcs2* WT), *Albumin-Cre*;*Hmgcs2*^fl/fl^ (or *Hmgcs2* LiKO), and *Villin-Cre*;*Hmgcs2^fl/fl^* (or *Hmgcs2* iKO) mice (12-32 weeks of age) were used. The *Hmgcs2* iKO mice were generated by crossing *Hmgcs2*^fl/fl^ mice with *Villin-Cre* mice (#004586, Jackson Laboratories, Maine, US); and the *Hmgcs2* LiKO mice were generated by crossing *Hmgcs2*^fl/fl^ mice with *Albumin-Cre* mice (#003574, Jackson Laboratories, Maine, US). All mouse strains were maintained on a C57BL/6J background and group-housed when possible in a 22 ± 2 °C temperature-controlled environment, kept on a 12:12 h light-dark cycle. Mice had *ad libitum* access to water and standard rodent chow diets (ssniff V1534-300, ssniff Spezialdiäten GmbH, Soest, Germany; Altromin no. 1310; Brogaarden, Denmark; or Inotiv Teklad, 2916) before the start of any experiments. Diet-induced obesity was induced by feeding young adult mice (>8 weeks old) a high-fat, high-sucrose diet (HFHSD, 5.16 kcal/g, S8022-E188, modified from D12492, ssniff Spezialdiäten GmbH, Soest, Germany) with a fat, protein, and carbohydrate content of 60.4% kcal, 17.2% kcal, and 22.4% kcal, respectively, including 18.4% sucrose by weight, for at least 12 weeks prior to interventions. Animal experiments on C57BL/6J and *Hmgcs2* LiKO mice were approved by the Government of Upper Bavaria, Germany, by the ethics committee of the State Office of Environment, Health and Consumer Protection of Brandenburg, Germany, performed according to institutional guidelines, and complied with the European Convention for the protection of vertebrate animals used for scientific purposes. Animal experiments on *Hmgcs2* iKO mice were performed in accordance with institutional regulations after protocol review and approval by Institutional Animal Care and Use Committees at the University of Texas Southwestern Medical Center.

### High-fat diets

Two high-fat diets (HFDs), LCFA HFD (4.89 kcal/g, D01060502G; Research Diets) and MCFA HFD (4.87 kcal/g, D18010905; Research Diets), were used in the chronic study. Both diets were comprised of 45% kcal fat, 35% kcal carbohydrate, and 20% kcal protein. The fat content contained 88% lard and 12% soybean oil in LCFA HFD, and pure MCT oil (60% C8:0 and 40% C10:0) in MCFA HFD. The energy content was determined by bomb calorimetry (IKA Werke, Germany) as previously described^21^.

### Oral gavage experiments

Double-housed, 1-4 h fasted chow-fed female and male mice (9-21 weeks of age), and male DIO mice (20-32 weeks of age) were randomized by body weight into 2, or 3 different groups and orally gavaged with 3-5 µL/g body weight of C8:0-MCT oil (glycerol trioctanoate, T9126, Sigma) or LCT oil (olive oil, O1514, Sigma, or lard, L0657, Sigma) or vehicle (drinking water). For experiments in indirect calorimetry cages, male DIO mice were single-housed and received 150 µL of C8:0-MCT or LCT oil (lard).

### Indirect calorimetry

Food intake, locomotor activity, energy expenditure, and substrate utilization were measured in 21-27-week-old DIO mice using a climate-controlled indirect calorimetric system (TSE Phenomaster System, Bad Homburg, Germany). Mice were randomly assigned to treatment groups and cage positions within the calorimetry system based on body weight. Animals were housed individually and allowed to acclimatize to the system for approximately 72 h before the experimental intervention. Water was orally administered on the two days prior to the intervention to facilitate habituation. On the experimental day, mice were fasted for 4 h (10.50 am - 2.50 pm) before receiving C8:0-MCT or LCT oil via oral gavage, with data acquisition starting at 3 pm. Metabolic parameters were continuously recorded for 24 h following treatment. Body weight was measured daily on a scale throughout the experiment.

### Blood glucose and ketone body measurements

Blood β-hydroxybutyrate (β-OHB) levels were measured by a ketone meter (FreeStyle Precision β-ketone, Abbott Laboratories, CH, IL, US; or GK+ Blood Glucose & Ketone Meter, Keto-Mojo, CA, US), and blood glucose concentrations were measured using a glucometer (CONTOUR NEXT, Bayer, Germany) with test strips on tail vein blood, portal vein blood, or systemic blood sampled before and at indicated time points after administration.

### Glucose Tolerance Test

Glucose tolerance test was performed after 6 h of fasting (6.30 am - 12.30 pm) with an intraperitoneal injection of D-glucose at 2 g/kg body weight in a volume of 10 mL/kg body weight. Blood glucose concentrations were measured in tail vein blood collected immediately before glucose administration and at 15, 30, 60, and 120 min thereafter using a handheld glucometer.

### Glucose-stimulated insulin secretion test

Glucose-stimulated insulin secretion test was commenced after a 6 h morning fast (6.30 am - 12.30 pm) with an oral gavage of D-glucose at 3 g/kg body weight in a volume of 10 mL/kg body weight. Blood glucose concentrations were measured in tail-vein blood collected immediately before glucose administration and at 2, 5, and 15 min thereafter using a handheld glucometer, with additional blood collected in EDTA-coated tubes at all time points for plasma insulin measurements.

### Portal vein blood collection

One hour post oral gavage, mice were sedated in a closed chamber with isoflurane. Sedation was maintained throughout with isoflurane. When the reflexes were absent, the portal vein was localized with an incision into the abdominal cavity and stabilized within tweezers. Portal vein blood samples were collected with a 27 G needle (with a syringe) inserted gently into the portal vein. Systemic blood was collected via cardiac puncture after portal vein blood collection, and mice were then euthanized by cervical dislocation.

### Tissue collection

All mice were killed with overdosed isoflurane or cervical dislocation. Tissues were immediately removed, and intestinal segments were further flushed with ice-cold phosphate-buffered saline (PBS) on ice. All tissues were snap-frozen in liquid nitrogen and stored at −80°C until further processing unless otherwise stated.

### Plasma analyses

Blood samples were collected in EDTA-coated tubes and immediately placed on ice until centrifugation (10 min at 1,000 x *g* at 4 °C). Plasma was stored at −80 °C until analysis. Plasma levels of insulin (Mouse Ultrasensitive Insulin ELISA, 80-INSMSU-E01, E10, Alpco, US), GDF15 (Mouse/Rat GDF-15 Quantikine ELISA Kit, MGD150, R&DSystems Inc.), and FGF21 (Mouse/Rat FGF-21 Quantikine ELISA Kit, MF2100, R&DSystems Inc.) were determined according to the manufacturer’s instructions.

### RNA Extraction and Quantification

RNA was isolated by Trizol-Chloroform extraction (peqGOLD TriFastTM, 30-2010P, VWR) from homogenized mouse tissues and precipitated with isopropanol. Genomic DNA was removed (DNAse I, EN0521, Fisher Scientific; RiboLock RNAse Inhibitor, EO0382, Fisher Scientific), and 1 µg RNA was transcribed into cDNA using LunaScript RT Super Mix (E3010L, New England Biolabs). Afterward, gene expression was quantified by qPCR using SYBR Green (Luna Universal qPCR Master Mix, M3003E, New England Biolabs) on a VIA7 Real-Time PCR System (4453534, Thermo Fischer). Primer efficiency was determined using an experiment-specific standard curve, and target gene expression was normalized to the expression levels of the appropriate housekeeping genes, *Hprt* and *Rpl13a*. The Primer sequences were as follows: *Hprt*: forward 5’-CTCATGGACTGATTATGGACAGGAC-3’, reverse 5’-GCAGGTCAGCAAAGAACTTATAGCC-3’; *Rpl13a*: forward 5’-GGAGGGGCAGGTTCTGGTAT-3’, reverse 5’-TGTTGATGCCTTCACAGCGT-3’; *Hmgcs2*: forward 5’-GAGGGCATAGATACCACCAACG-3’, reverse 5’-AATGTCACCACAGACCACCAGG-3’; *Acsm1*: forward 5’-AACATCCTCCCACCCAACAC-3’, reverse 5’-CATCATCGCCCCTTCCCAAG-3’; *Acsm3*: forward 5’-CCGTGGCAGCTAAATGTGAA-3’, reverse 5’-ACTTTGCCCAGCCTGTATCT-3’; *Acsm5*: forward 5’-GCTTCCGAGACTCCCAGATT-3’, reverse 5’-AGAAGCTTGGTTTGGAGGGA-3’.

### Western blotting

For Western blotting experiments, mice were fasted for 24 h prior to euthanasia and organ removal. Mouse tissues were homogenized with RIPA buffer (R3792, Teknova) supplemented with HALT protease and phosphatase inhibitors (78443, Thermo Fisher). Protein aliquots were loaded onto 4-20% Mini-PROTEAN TGX stain-free polyacrylamide gels (4568096, Bio-Rad) and transferred to PVDF membranes (Bio-Rad). The membranes were blocked in 5% milk in TBST for 30 min and incubated with the primary antibody, HMGCS2 (Abcam, ab137043, 1:8000 on the liver, and 1:1000 on the intestine and kidney) overnight at 4°C, followed by fluorescent secondary antibodies for one hour at room temperature. Bound antibodies were visualized using fluorescence and densitometry was determined based on total protein in Image Lab (Bio-Rad).

### Immunohistochemistry

For immunohistochemistry experiments, mice were fasted for 24 h prior to euthanasia and organ removal. Tissue samples were harvested and fixed in 10% neutral buffered formalin (SF100-4, Fisher Chemical) at 4°C overnight. Tissues were then embedded in OCT. Intestines were flushed with ice-cold PBS on ice prior to being fixed in cassettes, then jelly rolled and embedded in OCT. Eight µm sections were blocked in 5% normal donkey serum for one hour at room temperature. The sections were incubated overnight at 4°C with the primary antibody HMGCS2 (ab137043, Abcam, 1:150) and secondary antibodies for one hour at room temperature, DAPI for counterstain (62248, Thermo Fisher, 1:10,000). Slides were examined and images were obtained with the Keyence BZX-800 microscope and processed in ImageJ.

### Transcriptomic analysis on intestinal cells

Single-cell RNA-sequencing (scRNAseq) dataset of small intestinal (duodenum, jejunum, and ileum) crypts was extracted from GEO (GSE147319)^18^. Bulk RNAseq dataset of human jejunal crypts was extracted from GEO (GSE186509)^19^.

### Cell culture

#### Isolation and culture of primary hepatocytes

Primary murine hepatocytes were isolated from 12-16-week-old, chow-fed male C57BL/6J mice following a two-step collagenase perfusion protocol adapted from previous publications^42,43^. The mice were euthanized by isoflurane overdose, sterilized with 70% ethanol, and dissected to expose the inferior vena cava and portal vein. The inferior vena cava was then cannulated with a 25 G catheter needle and the liver was perfused using a peristaltic pump at 2-3 mL/min with Perfusion buffer I: PBS (without Ca^2+^/Mg^2+^) supplemented with 10 mM HEPES, 0.05% (w/v) KCl, 5 mM glucose, 0.2 mM EDTA, 0.001% (v/v) phenol red, at pH 7.4 and 37 °C. Once the liver started to blanch, the portal vein was cut to allow the blood to flow out. The perfusion was continued at ~6 mL/min for 5 min with Perfusion buffer I, followed by another 5 min perfusion with Perfusion buffer II: PBS (without Ca^2+^/Mg^2+^) supplemented with 10 mmol/L HEPES, 0.05% (w/v) KCl, 5 mmol/L glucose, 0.2 mmol/L EDTA, 1 mmol/L CaCl_2_, 0.001% phenol red, and 0.5 mg/mL collagenase IV (C5138, Sigma) at pH 7.4 and 37°C. After this two-step perfusion, the liver was removed, placed in a petri dish filled with ~10 mL perfusion buffer II, and torn with forceps. The crude hepatocyte solution was then filtered through 70 µm cell strainer, washed with 20 mL culture media: DMEM (PAN Biotech, P04-03596) supplemented with 10% (v/v) heat-inactivated FBS (P40-37500, Pan Biotech, heated at 55°C for 30 min), 1% (v/v) Penicillin/Streptomycin (15140122, Gibco), 0.05 nM insulin (I0516, Sigma), 100 nM Dexamethasone (D4902, Sigma), and 0.001 mg/mL Vitamin B_12_ (V2876, Sigma) followed by centrifugation at 50 *g* for 2 min at room temperature until the supernatant was clear. After the final wash, the primary hepatocyte pellets were resuspended in 25 mL culture media and the live cells were counted with trypan blue. The cells were seeded at a density of 6.25 x 10^4^/cm^2^ onto six-well plates that were previously coated with Collagen (1 mL 0.1 mg/mL Collagen I (CLS354236, Corning) diluted in 0.02N acetic acid). The cells were then incubated at 37 °C with 5% CO_2_ for 2-4 h and switched to serum-starvation culture medium for ~16 h until further treatment.

#### Culture of intestinal epithelial cell lines

Murine duodenal Mode-K cell line and colonic PTK-6 cell line^44^ were gifts from Dr. Sören Ocvirk (Gnotobiology, DIfE). Human colon cancer HT-29-MTX-E12 cell lines were purchased from Merck (12040401). The cells were seeded at 2 x 10^4^ cells/cm^2^ and cultured in DMEM supplemented with 5% FBS and 1% MEM NEAA (P08-32100, PAN Biotech) in a humidified incubator at 37 °C with 5% CO_2_ until reaching over 80% confluency.

#### Ketone body production experiment

Palmitic acid (PA) stock solution (100 mmol/L) was prepared by dissolving PA (P0500, Sigma) in 50% (v/v) ethanol and subsequently complexing it with 20% (w/v) bovine serum albumin (BSA, A7030, Sigma) at 37°C to reach a molar ratio of PA:BSA = 2:1. Octanoic acid (OA, C2875, Sigma) was dissolved in DMSO to prepare a 300 mmol/L OA stock solution.

On the day of the treatment, the media were replaced with freshly prepared media containing 300 µM OA, PA, or control. All wells contained 0.1% (v/v) DMSO, 0.15% (v/v) ethanol, and 1% (w/v) BSA. Six hours later, media were collected, and β-OHB concentrations were measured using Ketone Body Colorimetric Assay Kits (700190, Cayman) following the manufacturer’s protocol.

### Statistics

All bar graphs are presented as mean + standard error of the mean (SEM) with individual data points, and all time course data are presented as mean +/- SEM. Statistical analyses were performed using GraphPad PRISM 10.6.0 (GraphPad, CA, US) and are described in each figure legend. Post-hoc analysis was conducted when one-way ANOVA revealed an effect (*P* < 0.05); when two-way ANOVA revealed an interaction (*P* < 0.05) for experiments with two treatment groups; and when a significant main effect of diet was observed (*P* < 0.05), for experiments with three dietary groups. Statistical significance was defined as *P* < 0.05.

## Acknowledgments

We thank Dr. Sören Ocvirk for providing the intestinal cell lines. A.M.F. was funded by the Novo Nordisk Foundation (Grant number NNF22OC0074110) and the Independent Research fund Denmark (Grant number 10.46540/4253-00023B). C.C. is supported by the Novo Nordisk Foundation (Grant number NNF22OC0073778). The Novo Nordisk Foundation Center for Basic Metabolic Research is an independent Research Center, based at the University of Copenhagen, Denmark, and partially funded by an unconditional donation from the Novo Nordisk Foundation (www.cbmr.ku.dk) (Grant numbers NNF18CC0034900 and NNF23SA0084103). S.C.H was funded by National Institutes of Health (NIH) (Grant numbers R01DK135555, and R35GM137984) and an American Society of Nephrology Carl W. Gottschalk Research Scholar Grant. M.K. was supported by the Deutsche Forschungsgemeinschaft (DFG; KL 3285/5-1), the German Center for Diabetes Research (DZD; 82DZD03D03 and 82DZD03D1Y), the Novo Nordisk Foundation (Grant number NNF19OC0055192), and the Deutsche Diabetes Gesellschaft (DDG).

## Conflict of interest

C.C. is co-founder of Ousia Pharma ApS, a biotech company developing therapeutics for obesity. The remaining authors declare no competing interests.

## Notes

### Summary of Updates

New constitutive liver-specific Hmgcs2 knockout experiments, including studies in diet-induced obese mice, and additional acute and chronic metabolic phenotyping were added. Figures, results, methods, discussion, and supplementary Information were updated accordingly.

