## Supplemental Figures for "Metabolic benefits of dietary medium-chain fatty acids bypass hepatic ketogenesis"

### Fig. S1

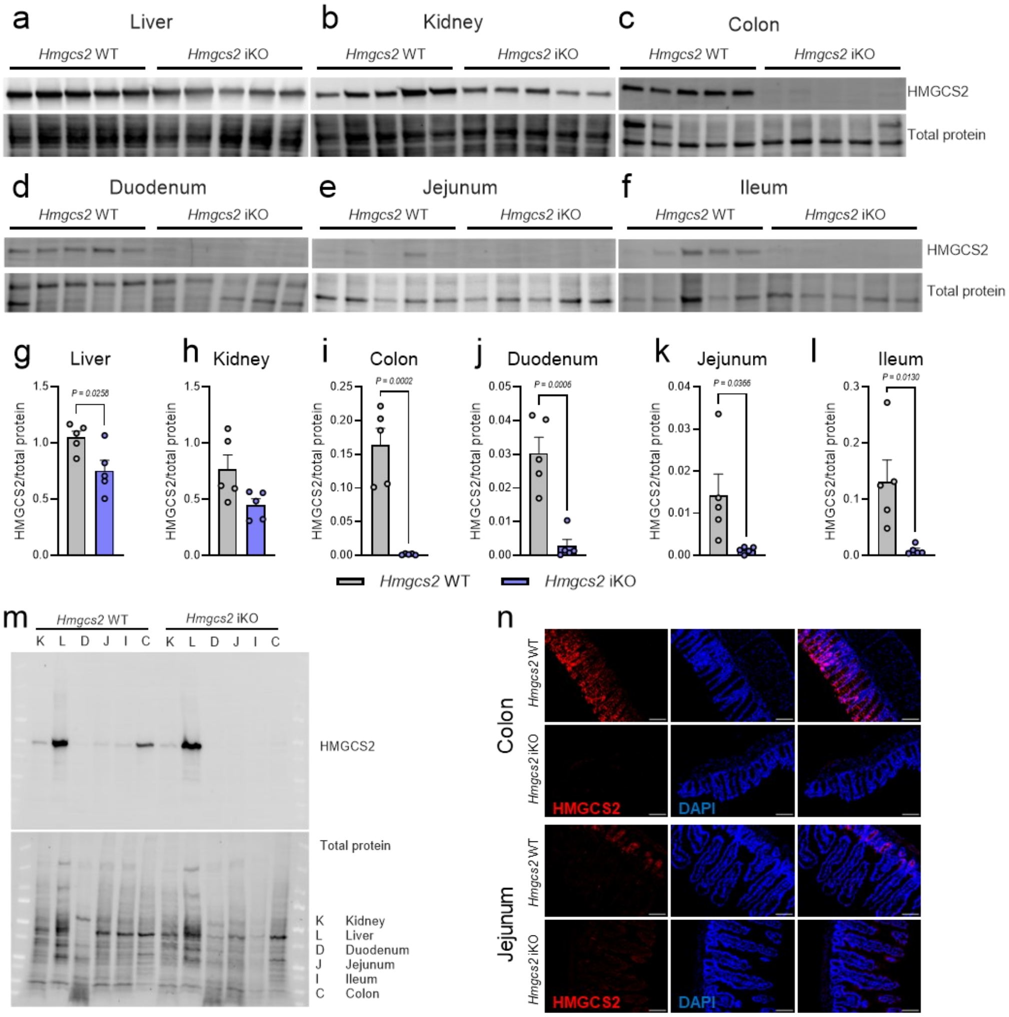


**Fig. S1 Intestinal 3-hydroxy-3-methylglutaryl-CoA synthase 2 (Hmgcs2) knockout validation**

a-l, representative western blots (a-f) and quantifications (g-l) of HMGCS2 protein in liver (a, g), kidney (b, h), colon (c, i), duodenum (d, j), jejunum (e, k), and ileum (f, l) from *Hmgcs2* iKO mice and their wild-type (WT) littermates in fasted state, n = 5. m, n, representative immunostaining images of colon (m) and jejunum (n) sections for HMGCS2 (red) and DAPI (blue) from *Hmgcs2* iKO and *Hmgcs2* WT mice in fasted state. Scale bars = 100 μM. Statistical significance was determined by unpaired t-tests (g-l). Data are shown as individual points with mean + SEM.

Fig S2

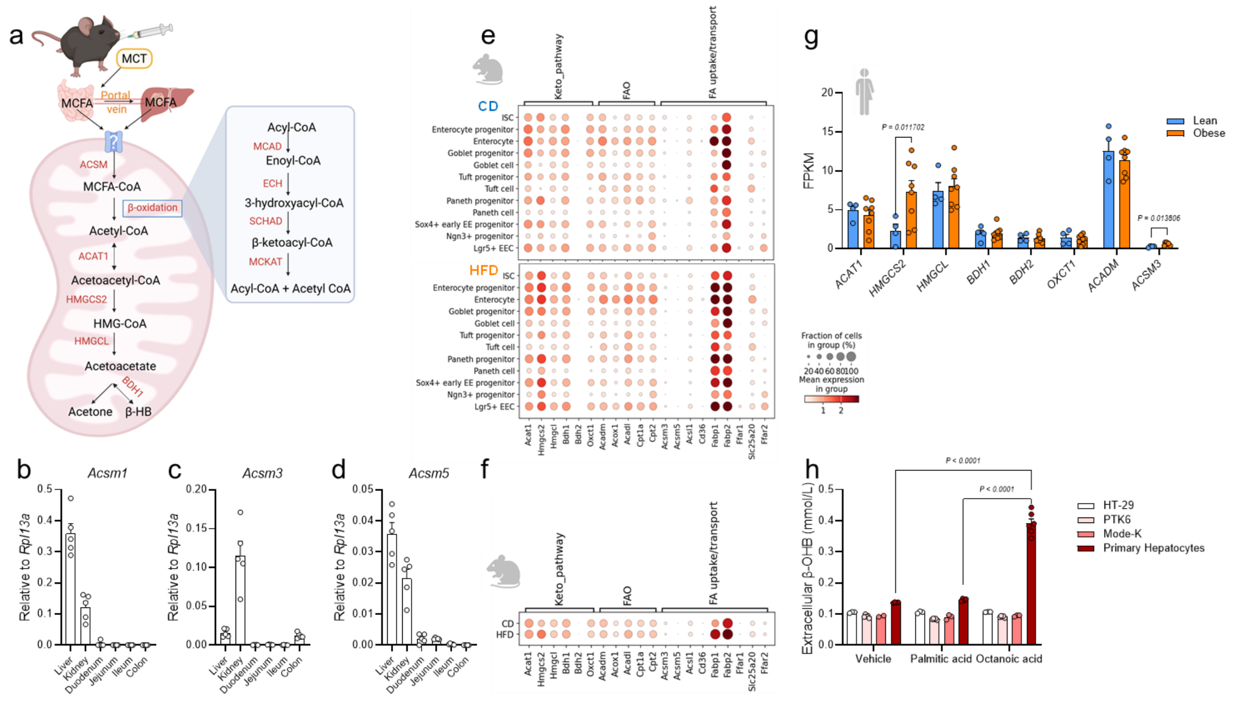


**Fig. S2 Intestines are not responsible for dietary MCT-induced hyperketonemia, likely due to the lack of medium-chain acyl-CoA synthetases**

a, Schematic of MCFA-induced hepatic ketogenesis pathway, created by BioRender.com. Ingested MCTs are transported from the intestines as MCFAs through the portal vein and enter the hepatic mitochondria, the primary site of ketogenesis. Medium-chain acyl-CoA synthetases (ACSMs) convert MCFAs into MCFA-CoA, which undergoes medium-chain acyl-CoA dehydrogenase (MCAD) initiated β-oxidation to produce Acetyl-CoA, the precursor of ketone bodies. b-d, mRNA abundance of *Acsm1* (b), *Acsm3* (c), and *Acsm5* (d) in the liver, kidney, duodenum, jejunum, ileum, and colon from male C57BL/6J mice, n = 5. e, f, transcriptomic analysis of jejunal crypts from mice fed on chow diet (CD) or high-fat diet (HFD); dot plot showing the expression of selected fatty acid metabolism related genes in small intestinal cells (e) or crypts (f), n = 3. g, transcriptional expression of selected fatty acid metabolism related genes in primary jejunal crypts from lean and obese subjects, n = 4-8. h, extracellular β-OHB concentrations in the media of intestinal cell lines and primary hepatocytes following 6 h incubation of vehicle, palmitic acid, or octanoic acid, n = 3-6. Data were analyzed by unpaired t-test (g) or two-way ANOVA with Tukey's post-hoc test (h). Data are shown as individual points with mean + SEM.

Fig S3

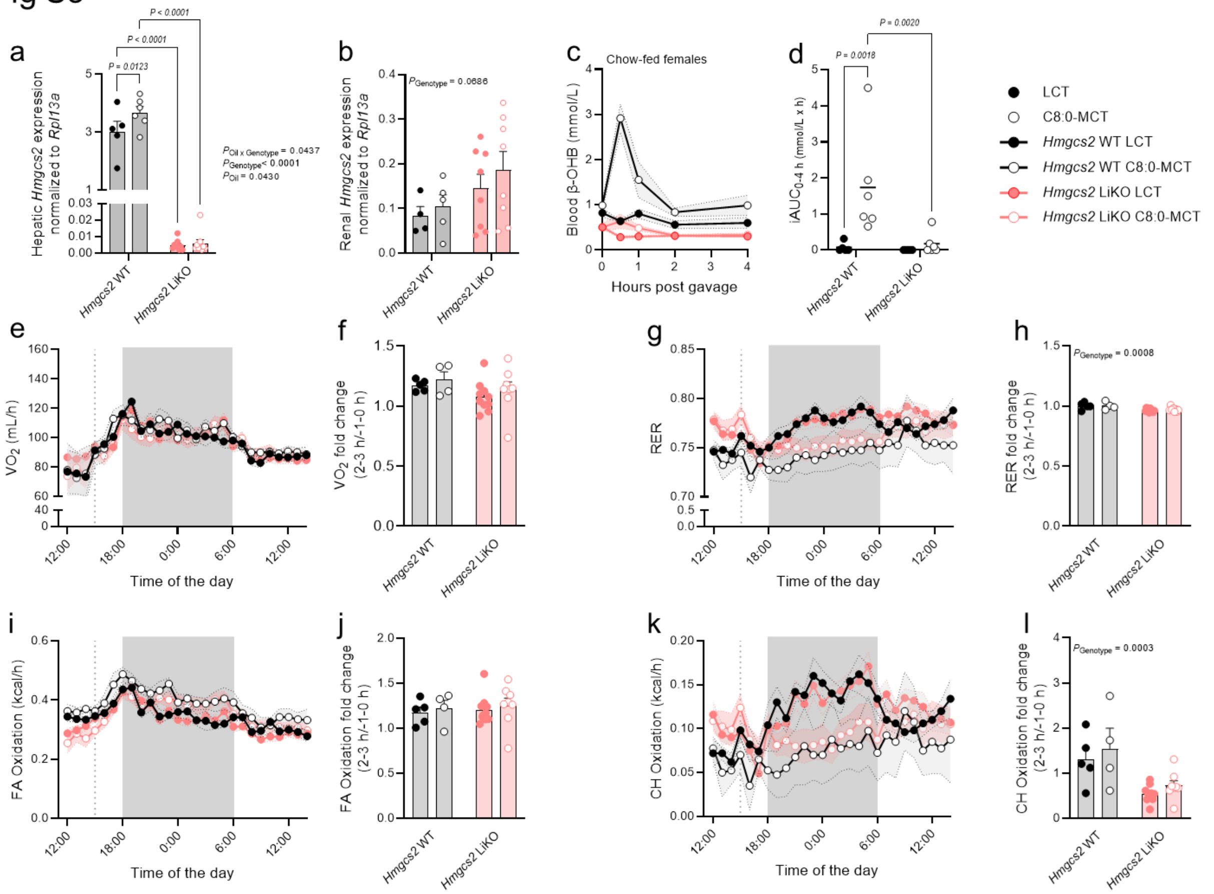


**Fig. S3 Blocking of hepatic ketogenesis lowers postprandial carbohydrate oxidation**

a, b, mRNA abundance of *3-hydroxy-3-methylglutaryl-CoA synthase 2* (*Hmgcs2*) in the liver (a), and kidney (b) of male DIO C57BL/6J mice, n = 5-9. c, d, chow-fed female C57BL/6J mice were gavaged with C8:0-MCT or LCT (Lard) control oil at 3 µL/g, followed by blood β-hydroxybutyrate (β-OHB) measurements at 0, 0.5, 1, 2, and 4 h (c), with incremental area under the curve (iAUC) (d), n = 5-6. e-l, metabolic phenotyping in indirect calorimetry cages after oral administration of C8:0-MCT or LCT at 150 µL per mouse in male DIO *Hmgcs2* WT or *Hmgcs2* LiKO mice, n = 4-10. e, f, oxygen consumption ($VO_2$) over 24 h (e), and paired pre- and early post-gavage $VO_2$, with lines connecting values from the same mouse; the early post-gavage window corresponds to 2-3 h after gavage (f). g, h, respiratory exchange ratio (RER) over 24 h (g) and paired pre-gavage and early post-gavage (h). i, j, fatty acid (FA) oxidation over 24 h (i) and paired pre-gavage and early post-gavage (j). k, l, carbohydrate (CH) oxidation over 24 h (k) and paired pre-gavage and early post-gavage (l). The dashed line indicates the time of gavage. Statistical significance was determined by two-way ANOVA with Fisher's LSD. Data are shown as mean +/- SEM (c, e, g, I, j, k) or individual points with mean + SEM (a, b, d, f, h, j, l).

Fig S4

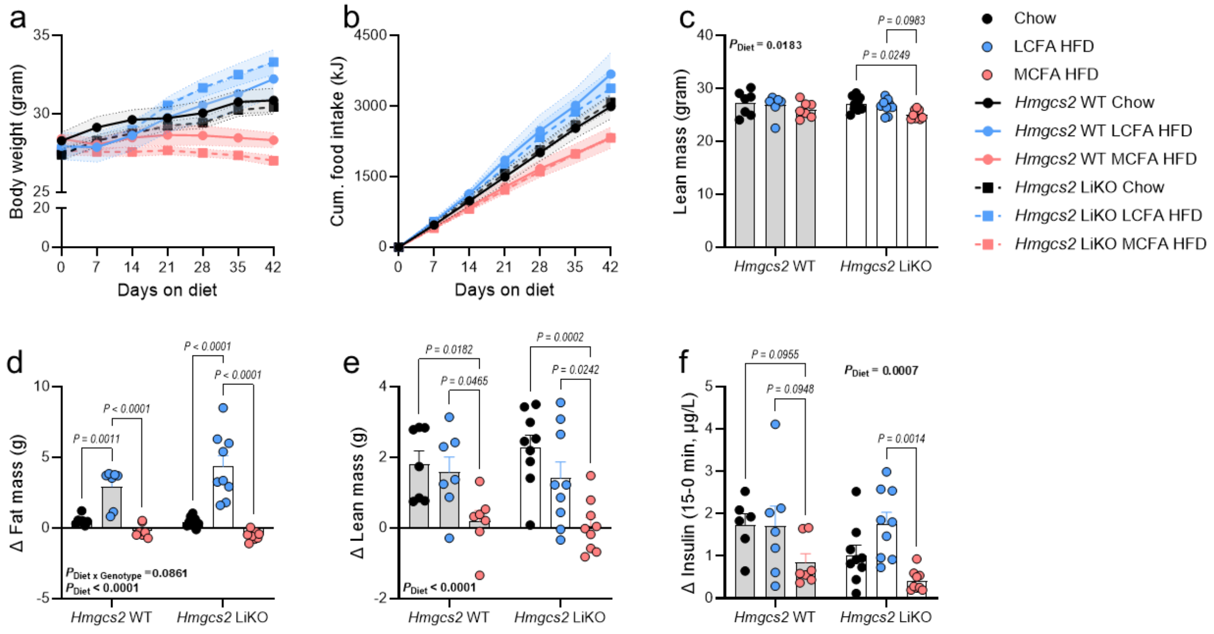


**Fig. S4 Dietary MCFAs reduce energy intake and fat mass independent of hepatic ketogenesis**

*Hmgcs2* WT or *Hmgcs2* LiKO mice were fed on the indicated diets for 6 weeks. a, b, body weight (a, n = 7-9), and food intake (b, n = 3-4) were recorded at designated time points. c-e, lean (c), and fat mass (d) were determined pre-diet switch, and at week 6 post-diet switch; changes in fat mass (d) and lean mass (e) were calculated; n = 7-9. f, 15-min insulin secretion changes determined during an oral glucose stimulated insulin secretion test (3 g of glucose per kg body weight) conducted 5 weeks after placing mice on the indicated diets. f, plasma FGF21 concentrations at 6 weeks after putting mice on indicated diets. Statistical significance was determined by two-way ANOVA with Tukey's post-hoc test. Data are shown as mean +/- SEM (a, b) or individual points with mean + SEM (c-f).
